# NoisyFlow: Differentially Private Optimal Transport Using Neural Networks for Secure Biomedical Data Sharing

**DOI:** 10.1101/2025.01.31.635830

**Authors:** Yunyang Li, Nikhil Khandekar, Skylar Wang, Varada Khanna, Julian Sanker, Mark B. Gerstein

## Abstract

**Motivation:** Advancing data sharing in biomedical research, particularly for sensitive genomic and clinical datasets, is crucial for improving model performance across diverse patient populations. However, stringent privacy concerns hinder collaboration and limit insights derived from multi-institutional datasets. Current approaches to privacy-preserving data sharing fail to address gaps between data distributions.

**Results:** We introduce NoisyFlow, a differentially private neural network-based optimal transport framework designed to enable secure and unbiased biomedical data sharing. By integrating optimal transport theory with neural networks and differential privacy mechanisms, our framework aligns data distributions across institutions while preserving individual privacy. NoisyFlow eliminates the need for direct data sharing and reduces distribution shifts caused by covariate and batch effects. Empirical evaluations demonstrate the framework’s effectiveness in handling high-dimensional single-cell genomic data and histopathology images, achieving superior privacy guarantees while maintaining high utility in downstream tasks such as disease classification.

**Availability and implementation:** The implementation of NoisyFlow is available at https://github.com/liyy2/NoisyFlow.

**Supplementary information:** Supplementary data are available online.

## 1 Introduction

### High level description of the problem

Recent advances in precision medicine and biomedical research have led to the aggregation of extensive patient data sets, providing substantial opportunities to elucidate disease mechanisms and develop targeted therapies (Roberts *et al*., 2020; Chen *et al*., 2020). For example, large-scale genomic analyzes can identify biomarkers associated with complex diseases, while comprehensive data sets of electronic health records (EHR) can facilitate the discovery of previously unrecognized risk factors. However, effectively leveraging these rich data sources requires addressing two interrelated challenges. The first challenge is ensuring patient privacy while sharing sensitive medical data across institutions (Wang *et al*., 2023; Jwa and Poldrack, 2022). The second challenge involves managing the heterogeneity of biomedical datasets, particularly in mitigating distribution shifts that arise when integrating populations from diverse clinical settings. Such challenges have led to pervasive data silos that fragment information and limit cross-institutional collaboration (Wan *et al*., 2022; Alrefaei *et al*., 2022; Amorim *et al*., 2022; Bonomi *et al*., 2020; Clayton *et al*., 2019). While enhanced data sharing could substantially improve machine learning model performance by exposing algorithms to larger and more diverse patient populations, privacy concerns often preclude such cooperation. These concerns are well-founded, as breaches in biomedical data privacy can potentially lead to insurance discrimination, stigmatization, or employment prejudice (Wan *et al*., 2022; Alrefaei *et al*., 2022; Amorim *et al*., 2022; Bonomi *et al*., 2020; Clayton *et al*., 2019). Consequently, models are typically trained on homogeneous datasets, which are vulnerable to algorithmic bias and generally exhibit poor performance on out-of-distribution (OOD) populations (Zou and Schiebinger, 2021). This limitation can potentially result in misdiagnoses and suboptimal treatment strategies.

### Differential Privacy in Biomedical Data

Addressing data-sharing challenges in the biomedical domain necessitates robust privacy-preserving approaches. Differential privacy (DP) has emerged as a prominent solution, offering mathematically rigorous guarantees that an algorithm’s output remains statistically indistinguishable whether or not a specific individual’s data is included. In essence, DP ensures that no single patient’s information can be inferred from a model’s outputs, even if a malicious actor has access to auxiliary datasets or prior knowledge. This property can be particularly valuable when dealing with high-dimensional datasets such as genomics or medical imaging, where the risk of re-identification is nontrivial.

### Distribution Shifts and Optimal Transport

Simultaneously, the challenge of aligning data distributions from disparate sources necessitates methods that can preserve the intrinsic structure of these datasets. Consider two healthcare centers—one in Boston and another in Tokyo. The Boston hospital may collect MRI scans with different imaging protocols and patient demographics compared to the Tokyo healthcare center, resulting in datasets with substantially distinct statistical and clinical properties. Even after ensuring privacy through DP-based federated learning, the combined dataset might exhibit shifts in patient ethnicity, age, or comorbidities, leading to differing underlying probability distributions (Alhamoud *et al*., 2024). Naively merging these data could degrade model performance or obscure important subgroup-specific patterns.

Optimal transport (OT) is particularly effective in bridging these gaps. OT seeks the most efficient transformation of one probability distribution into another, subject to constraints that preserve key structural relationships in the data. When applied to biomedical datasets, OT can align data from different populations in a geometry-aware manner, considering local relationships, cluster structures, and other distributional properties (Zhang *et al*., 2021). This is especially advantageous in complex, multimodal biomedical settings where patients or conditions naturally cluster into separate but related subpopulations (Zhang *et al*., 2021; Cao *et al*., 2021).

Here, we present NoisyFlow, a framework that integrates differential privacy with optimal transport (OT) to facilitate secure and efficient knowledge transfer between institutions. By preserving privacy and improving geometry-aware data alignment, NoisyFlow offers a promising approach for collaborative biomedical research. We include a visual representation of NoisyFlow in Figure 1. Its implementation has the potential to support the development of collaborative research while adhering to stringent patient privacy standards.

**Fig. 1.**
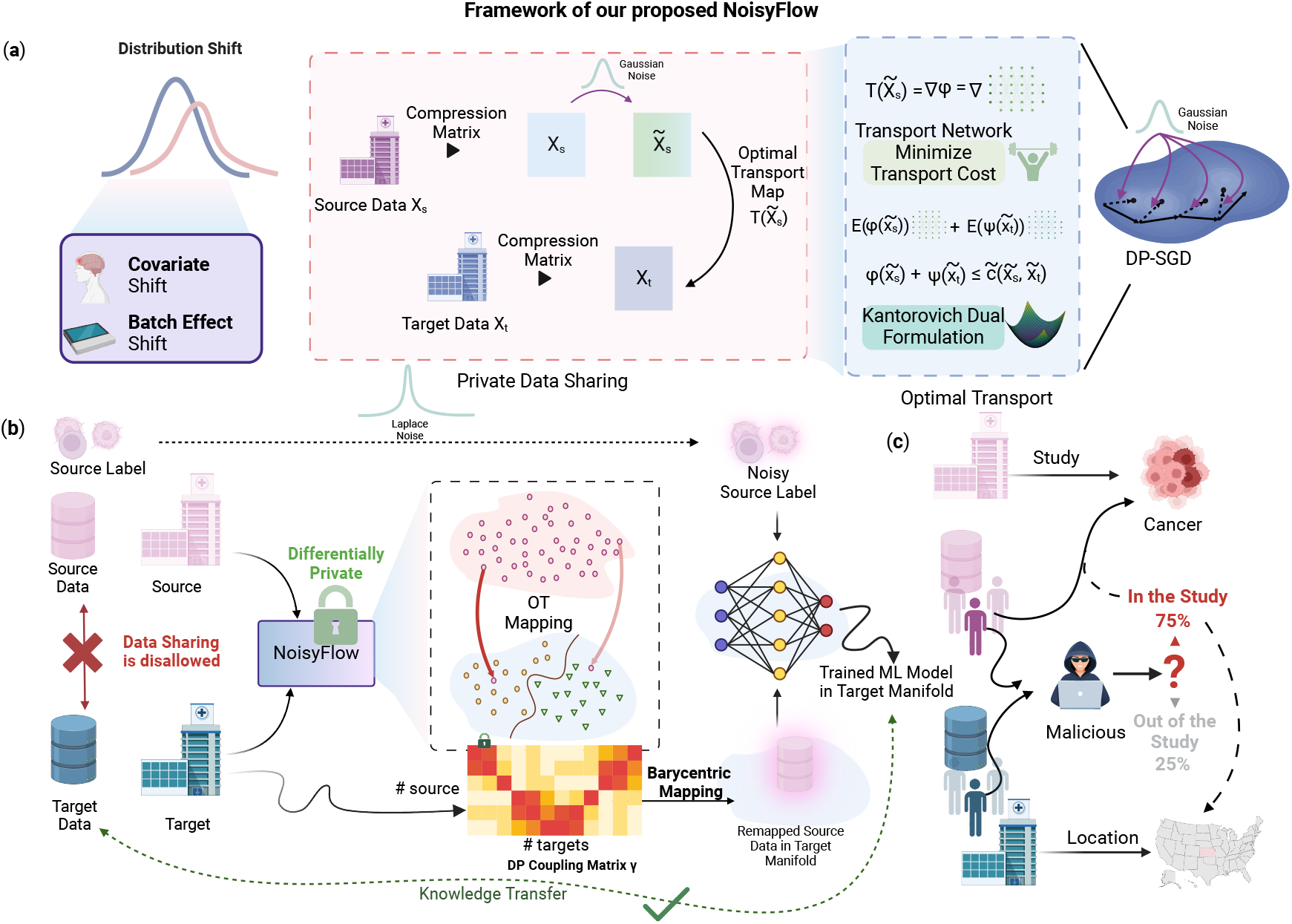
Overview of the NoisyFlow framework for differentially private knowledge transfer under distribution shifts. (a) Underlying motivation for addressing covariate shift and batch effect shift, depicted by the mismatched distributions of source and target data. Both source *X*_*s*_ and target *X*_*t*_ are compressed via compression matrices to produce low-dimensional representations 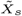 and 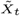. Additive Gaussian noise is introduced for privacy protection before computing the optimal transport mapping 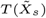, which is formulated through the Kantorovich dual problem to minimize transport cost. The final transport map *T* is parameterized by potentials *φ* and *ψ*, whose gradients transfer source data into the target manifold under differential privacy constraints (DP-SGD). (b) Differential privacy is enforced during cross-institutional data transfer. While the source data and labels cannot be directly shared, NoisyFlow allows knowledge transfer from the source to the target domain by perturbing labels (Laplace noise) and learning a private OT coupling matrix *γ*. This matrix aligns source and target samples in a unified manifold, enabling barycentric mapping and preserving task-related structure. The colored heatmap depicts the learned coupling between source and target populations, capturing correspondences in a privacy-preserving manner. Once mapped onto the target manifold, a downstream machine learning (ML) model (e.g., a neural network) is trained with the remapped source data and noisy source labels. (c) NoisyFlow preserves utility for clinical tasks, such as cancer classification, while controlling disclosure risks. The schematic on the right highlights potential membership inference concerns, indicating how malicious entities might leverage study participation to determine sensitive information. The NoisyFlow framework mitigates these risks through strict privacy guarantees conferred by differential privacy at every stage of the knowledge transfer.

### Contributions

The contributions of our work can be summarized as follows:

- **Conceptual Innovation:** We explore the application of differentially private neural-network based optimal transport (NoisyFlow) to enhance the privacy of genomics data. By integrating differential privacy with optimal transport, we propose an approach to address the privacy-utility trade-off in cross-institutional collaborative research, facilitating secure knowledge transfer without direct data sharing.
- **Technical Development:** We provide an analysis of the proposed NoisyFlow framework, including an examination of its theoretical properties and an approximation error analysis. This analysis aims to quantify the trade-offs between privacy guarantees and data utility.
- **Empirical Evaluation:** We conduct extensive empirical evaluations on real-world datasets, with a particular focus on single-cell genomics data and medical imaging as detailed in the dataset section. Our experiments suggest that the proposed approach effectively preserves privacy while maintaining utility in downstream tasks, such as disease classification.

### Related Prior Works

As biomedical datasets grow in complexity and volume, privacy challenges are becoming increasingly critical. Biomedical research encompasses a wide range of data modalities, including functional genomics (Gürsoy *et al*., 2022) and medical imaging, such as chest X-rays (Packhäuser *et al*., 2022). In particular, large-scale imaging datasets have driven significant advancements in cancer detection, such as for skin and breast cancer (M., 2019; Litjens *et al*., 2018). However, the widespread availability and reuse of such datasets introduce substantial privacy risks, including direct genotyping from raw sequencing reads (Pajuste *et al*., 2017), inference of genotypes from expression values (Schadt *et al*., 2012), and exposure of sensitive phenotypic information through imaging analyses (Panzade *et al*., 2024). For example, expression quantitative trait loci (eQTLs) can reveal individual genotypes (Harmanci and Gerstein, 2016), and imaging data might inadvertently contain identifiable health conditions. Addressing these challenges requires innovative privacy-preserving methods tailored to the multifaceted nature of biomedical data. Current approaches like privacy-preserving BAM files (Gürsoy *et al*., 2020), homomorphic encryption (Gentry, 2009), and blockchain-based systems (Warnat-Herresthal *et al*., 2021) aim to balance privacy protection with data utility. However, these methods often face scalability issues as biomedical datasets continue to grow in size and complexity. Additionally, the transient nature of certain data types means that privacy risks can evolve over time, requiring dynamic privacy frameworks to effectively safeguard sensitive information. Ensuring robust and scalable privacy measures is essential for the responsible advancement of biomedical research.

Further, recent advancements in differential privacy have led to the development of robust frameworks designed to protect biomedical data while maintaining its research utility. One key challenge in this domain is handling distribution shifts in biomedical datasets, which can affect the generalizability of models. Koh *et al*. (2021) introduced benchmarks for studying such shifts, including those present in widely used datasets like CAMELYON for breast cancer detection (Litjens *et al*., 2018). To address these challenges, LeTien *et al*. (2019) proposed a differentially private optimal transport framework for domain adaptation, demonstrating its effectiveness in mitigating distribution shifts while preserving privacy.

Building on this, DP-Sinkhorn was proposed as an optimal transport-based method that avoids the instability issues commonly associated with GAN-based approaches while preserving differential privacy (Cao *et al*., 2021). This method has proven particularly effective for high-dimensional data synthesis, such as generating informative RGB images under strict privacy constraints without requiring auxiliary public data.

In the realm of genomic data, specialized frameworks incorporating local differential privacy have been developed to address the unique challenges posed by correlated genetic information (Yilmaz *et al*., 2021). He *et al*. (2018) focused primarily on differential privacy for genomic data releases, highlighting limitations in handling complex genetic correlations (He *et al*., 2018). Privatedriver exemplifies a privacy-preserving system that identifies cancer subtype-specific driver genes from multigenomics data, but it generally lacks comprehensive privacy guarantees (Song *et al*., 2024).

Optimal transport has been applied to patient-level distance detection in single-cell genomic data, offering a promising approach for privacy integration. However, it continues to face challenges in scalability and accuracy (Joodaki *et al*., 2023). Bunne *et al*. (2023) introduced CellOT, leveraging optimal transport theory within neural network architectures to map complex high-dimensional distributions, but this model is not scalable to high-dimensional data (Bunne *et al*., 2023). Recent studies like TIGON have addressed the scalability of high-dimensional data by managing large-scale trajectory inference using unbalanced optimal transport (Sha *et al*., 2023).

## 2 Preliminaries / Problem Statement

We consider a general setting where 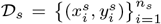 denote the labeled data from the *source* domain, and 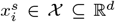 (e.g., RNA expressions of different genes or a set of lab test results) and 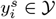 (e.g., clinical outcomes or phenotypic traits). Similarly, let 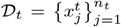 denote the unlabeled data from the *target* domain, which may represent a different institution, experimental condition, or demographic group. Our primary goal is to leverage knowledge from the labeled source domain 𝒟_*s*_ to make accurate predictions for the unlabeled target data 𝒟_*t*_, thereby enabling effective knowledge transfer. Here, we assume a general setting without loss of generality, and our framework accommodates various scenarios, including cases where the target domain is also labeled.

A key challenge arises when the source and target domains follow different distributions, typically referred to as a *distribution shift*. Concretely, let *µ*_*s*_ and *µ*_*t*_ be the probability measures (distributions) for the source and target domains. The distribution shift may occur due to various factors, including *batch effects* and *covariate shifts*.

### Definition 2.1

(Batch Effects). *Batch effects represent a distributional shift in observed data, arising from technical artifacts introduced by varying experimental or collection conditions across different batches. These effects cause systematic deviations that can obscure the true underlying biological (or statistical) signal*.

### Definition 2.2

(Covariate Shift). *Covariate shift arises when the distribution of input features differs between the source and target domains, while the conditional distribution of labels given those features remains the same. Formally, let* (*x, y*) *denote a feature-label pair sampled from a distribution p*_*s*_(*x, y*) *in the source domain and p*_*t*_(*x, y*) *in the target domain. Covariate shift means p*_*s*_(*y* | *x*) = *p*_*t*_(*y* | *x*) *but p*_*s*_(*x*) ≠ *p*_*t*_(*x*). *For example, a diagnostic model trained on a younger patient population may face covariate shift when deployed for older patients if the relationship between features and clinical outcomes (i*.*e*., *p*(*y* | *x*)*) remains unchanged, but the distribution of patient features (e*.*g*., *comorbidities, genetic backgrounds) differs across age groups*.

Understanding and addressing these types of distribution shifts are crucial for developing robust machine learning models that generalize well across diverse biomedical datasets.

### 2.1 Optimal Transport Framework

Optimal transport (OT) provides a principled framework for comparing and aligning distributions *µ*_*s*_ and *µ*_*t*_. By interpreting the source and target as probability measures on 𝒳, OT finds a coupling (or a mapping) that *minimizes* the total cost of transporting mass from *µ*_*s*_ to match *µ*_*t*_.

#### p-Wasserstein Distance

To quantify the distance between these probability measures, a commonly used OT distance is the *p*-Wasserstein distance, defined as follows.

##### Definition 2.3

(*p*-Wasserstein Distance). *Let µ*_*s*_, *µ*_*t*_ ∈ 𝒫 (𝒳) *be two probability measures on* 𝒳. *The p*-Wasserstein distance *between µ*_*s*_ *and µ*_*t*_ *is defined by*

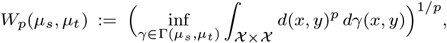

*where d*(*x, y*) *is a distance function*, Γ(*µ*_*s*_, *µ*_*t*_) *denotes the set of all couplings (joint probability measures) on* 𝒳 *×* 𝒳 *whose marginals are µ*_*s*_ *and µ*_*t*_.

Building upon this foundational distance metric, we delve deeper into the optimization framework that underpins OT.

##### Definition 2.4

(Primal OT and Optimal Coupling). *Let µ*_*s*_ *and µ*_*t*_ *be probability measures on a Polish space 𝒳, and let c* : 𝒳 *×* 𝒳 → [0, ∞) *be a given cost function. The* primal optimal transport problem *seeks a coupling*

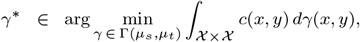

*where* Γ(*µ*_*s*_, *µ*_*t*_) *is the set of all couplings, that is, all joint probability measures γ on* 𝒳 × 𝒳 *whose marginals are µ*_*s*_ *and µ*_*t*_. *We refer to any such minimizer γ*^*^ *as the* optimal coupling *or* optimal transport plan.

In practice, *µ*_*s*_ and *µ*_*t*_ are often *empirical* measures derived from finite samples. Namely,

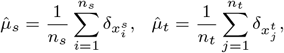

where *δ*_*x*_ is the Dirac measure at *x*. The discrete OT problem becomes:

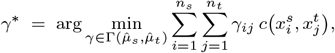

subject to

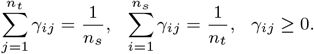

##### Definition 2.5

(Kantorovich Dual Problem). *To gain further insights into the structure of the OT problem, we explore its dual formulation*.

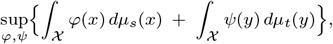

*subject to the constraint ϕ*(*x*) + *ψ*(*y*) ≤ *c*(*x, y*) *for all x, y* ∈ 𝒳. *Here, ϕ and ψ are called the* dual potentials.

Under appropriate regularity conditions on the cost function *c*, strong duality holds, ensuring that the optimal values of the primal and dual problems are equal. This result connects the primal and dual formulations of OT. We now introduce a fundamental theorem that characterizes the optimal transport map under specific conditions.

##### Theorem 2.1

(Brenier’s Theorem). *Let µ*_*s*_ *and µ*_*t*_ *be two probability measures on* ℝ^*d*^ *such that µ*_*s*_ *is absolutely continuous with respect to the Lebesgue measure. Then there exists a unique optimal transport plan γ*^*^ *between µ*_*s*_ *and µ*_*t*_, *and moreover, this plan is induced by a deterministic map T* ^*^ : ℝ^*d*^ → *ℝ*^*d*^. *In fact, there is a convex function* Φ : ℝ^*d*^ → ℝ *such that*

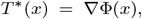

*and the pushforward of µ*_*s*_ *by T* ^*^ *equals µ*_*t*_. *Equivalently, the unique solution γ*^*^ *is supported on the set {*(*x*, ∇Φ(*x*)) : *x* ∈ ℝ^*d*^*}*.

Here, *T* ^*^ : *𝒳* → *𝒳*, is called the *optimal transport map*. For the quadratic cost 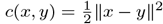, Brenier’s theorem establishes that *T* ^*^ is the gradient of a convex potential function *φ*: *T* ^*^(*x*) = ∇*φ*(*x*),. Formally, this is given as:

##### Lemma 2.2

(Optimal Transport Map from Dual Potentials). *Let µ*_*s*_ *and µ*_*t*_ *be probability measures on* ℝ^*d*^, *and let φ, ψ be the optimal dual potentials satisfying*

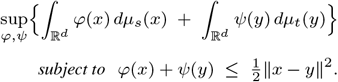

*If µ*_*s*_ *is absolutely continuous, then one can take*

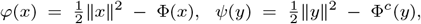

*where* Φ *is a convex function and* Φ^*c*^ *is its c-transform. Moreover, the optimal transport map is*

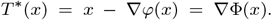

##### Remark 2.3

(Relationship of the Optimal Transport Map and the Optimal Coupling) *In many settings, especially with the squared-Euclidean cost* 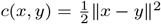, *the optimal coupling γ*^*^ *is further induced by a* deterministic *map T* ^*^ : 𝒳 → 𝒳. *That is, there exists a function T* ^*^ *such that*

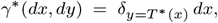

*meaning mass at x is sent* entirely *to the point T* ^*^(*x*). *In this situation, T* ^*^ *is called the* optimal transport map, *or the* Monge solution. *However, for more general costs or measure types, the optimal coupling may* not *be induced by a single map; the solution can be a more general joint distribution. Throughout, we may refer interchangeably to γ*^*^ *as the* optimal transport plan.

To further establish the connection between the optimal transport map and the coupling matrix, we introduce the concept of barycentric mapping:

##### Definition 2.6

(Barycentric Mapping). *Let µ*_*s*_ *and µ*_*t*_ *be two probability measures on a Polish space 𝒳, and let γ*^*^ *be an optimal coupling (or transport plan) between them. The* barycentric mapping *induced by γ*^*^ *is the map T*_*γ*_ : 𝒳 → 𝒳 *defined by*

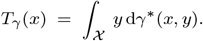

In other words, for each point *x* in the support of the source measure *µ*_*s*_, *T*_*γ*_ (*x*) is the barycenter (or center of mass) of all points *y* in the target domain, weighted by *γ*^*^(*x, y*). Hence, *T*_*γ*_ maps each source point to the average location in the target distribution under the optimal transport plan.

### 2.2 Neural Network-Based Approximation of Dual Potentials

In high-dimensional settings, the dual potentials *φ* and *ψ* can be efficiently approximated using neural networks. Let *φ*_*θ*_ and *ψ*_*φ*_ represent neural networks parameterized by *θ* and *φ*, respectively. The dual problem becomes:

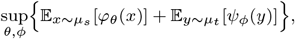

subject to *φ*_*θ*_(*x*) + *ψ*_*φ*_(*y*) ≤ *c*(*x, y*) for all *x, y*.

For the quadratic cost 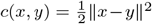, the optimal transport map *T* ^*^ can be parameterized as the gradient of a convex neural network *ψ*_*θ*_. Using input-convex neural networks (ICNNs), which ensure the convexity of *ψ*_*θ*_, the transport map is given by: *T* ^*^(*x*) = ∇*ψ*_*θ*_(*x*).

## 3 Differential Private Distribution Alignment

In the previously defined setting, distribution alignment involves learning a mapping or transport plan to align two probability measures: the source distribution *µ*_*s*_ and the target distribution *µ*_*t*_. Given the sensitive nature of the data in applications such as healthcare, where the source and target datasets often contain personal or identifiable information, it is essential to ensure that the learned alignment respects the privacy of individual data points. To achieve this, we introduce a notion of *differentially private distribution alignment*, which guarantees that no adversary can infer sensitive details about any single individual in both datasets. Formally, we formulate this as:

### Definition 3.1

(DP Distribution Alignment). *An alignment algorithm 𝒜*(𝒟 _*s*_, 𝒟 _*t*_) *that outputs a transport plan T* : ℝ^*d*^ → ℝ^*d*^ *is said to be* (*ε*, δ)*-differentially private if, for any pair of neighboring source datasets* 𝒟_*s*_ *and* 𝒟′_*s*_ *(differing by at most one data point), and for any measurable set 𝒯 in the space of possible transport plans, we have*

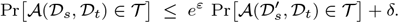

In this context, the privacy budget *ε >* 0 governs the balance between privacy and accuracy, with lower values enforcing more stringent privacy protections. The parameter δ ≥ 0 permits a minimal probability of privacy guarantee breaches, offering flexibility in settings where absolute privacy is infeasible. The algorithm 𝒜 denotes the computational procedure that transforms one dataset into another via the transport mechanism. The fundamental equation delineates the relationship between the probability that 𝒜 yields an output within a designated set 𝒯 when applied to dataset 𝒟_*s*_, versus the probability of producing the same output when the input dataset is slightly modified to 𝒟′_*s*_. The term *e*^*ε*^ modulates this probability disparity, with *ε* determining the extent of permissible privacy loss, while *δ* accommodates a negligible margin of error. This framework ensures that the algorithm’s behavior remains consistent, preventing its outputs from disclosing substantial information about individual data points and thereby maintaining privacy.

## 4 The model/framework

In differential privacy, the scale of additive noise typically increases with the “sensitivity” of the mechanism—i.e., the extent to which small changes in the input dataset can affect the output. In high-dimensional spaces, norms (and consequently distances) can grow large or vary significantly. Projecting the data to a lower-dimensional space (*k ≪ d*) helps standardize and often reduce the worst-case sensitivity. To achieve this, we apply the Johnson-Lindenstrauss (JL) transform. This involves generating a random matrix 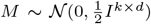 and projecting the data into a lower-dimensional space of dimension *k*. The JL transform preserves the Wasserstein distance of the original data up to a constant factor. Specifically, the source data is transformed as 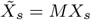, and the target data is transformed analogously as 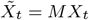.

To preserve differential privacy, we introduce Gaussian noise to the transformed source data, resulting in noisy transformed data 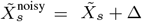, where 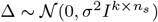 represents the added noise. This ensures that the source data are not revealed during the transport process, satisfying the differential privacy constraints.

### Bias Correction

The introduction of noise to the source data increases the expected cost between data points. Intuitively, noise inflates the uncertainty in the distribution, increasing its entropy and thus resulting in a higher cost To correct for this effect, we adjust the cost function by subtracting a term proportional to the noise variance. Specifically, we modify the original cost function *c*(*x*_*s*_, *x*_*t*_) (often chosen as the squared Euclidean distance) to obtain an adjusted cost function:

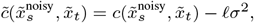

where *𝓁 >* 0 is a scaling parameter, and 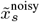 and 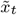 are the noisy transformed source data and transformed target data, respectively. This adjustment ensures that the expected cost remains unbiased despite the noise added to the source data.

#### Lemma 4.1

(Unbiasedness of Adjusted Cost). *Suppose c*(*u, v*) = | | *u* − *v* | | ^2^, *and let* 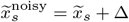 *with* Δ ∼ 𝒩 (0, *σ*^2^*I*_*k*_). *Then*

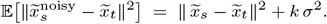

*Consequently*,

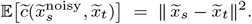

Proof Sketch. Expand the squared norm:

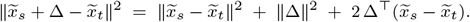

Taking expectation over Δ ∼ 𝒩 (0, *σ*^2^*I*_*k*_) yields 𝔼[| | Δ | | ^2^] = *k σ*^2^ and 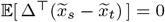. Hence the result follows.

To implement the optimal transport plan, we adopted the dual formulation. In order to make the dual formulation computationally feasible and scalable to high-dimensional data, we parameterize the dual potentials *ϕ* and *ψ* using neural networks. Let *ϕ*_*θ*_ : ℝ ^*k*^ → ℝ and *ψ*_*ϕ*_ : ℝ ^*k*^ → ℝ be neural networks with parameters *θ* and *η*, respectively, that map the noisy transformed source data 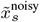 and transformed target data 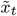 to scalar potentials. This neural network-based parameterization enables the model to learn complex relationships between the data points in the transformed space while leveraging the flexibility of deep learning.

The optimization objective in the dual form of the problem is to minimize the following loss function:

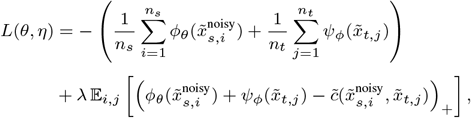

where *λ >* 0 is a penalty parameter that enforces the constraint 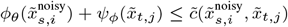, and (·)_+_ denotes the positive part function. The term 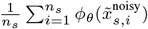 represents the expectation of the source potential over the noisy transformed source data, and similarly for the target potential.

### 4.1 Algorithm for NoisyFlow

Here, we outlined the formulation of NoisyFlow in Algorithm 1. Specifically, the adjusted cost matrix 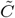 is computed by subtracting the noise adjustment term *𝓁σ*^2^ from the original cost function, which in our case is the squared Euclidean distance. The neural networks *ϕ*_*θ*_ and *ψ*_*ϕ*_ are updated iteratively using *differentially private stochastic gradient descent* (DP-SGD) to minimize the loss function, while ensuring that the constraints on the dual potentials are enforced through the penalty term. DP-SGD also protects the privacy of the target domain before target domain publishes the optimal transport plan to the source.

#### Algorithm 1

NoisyFlow

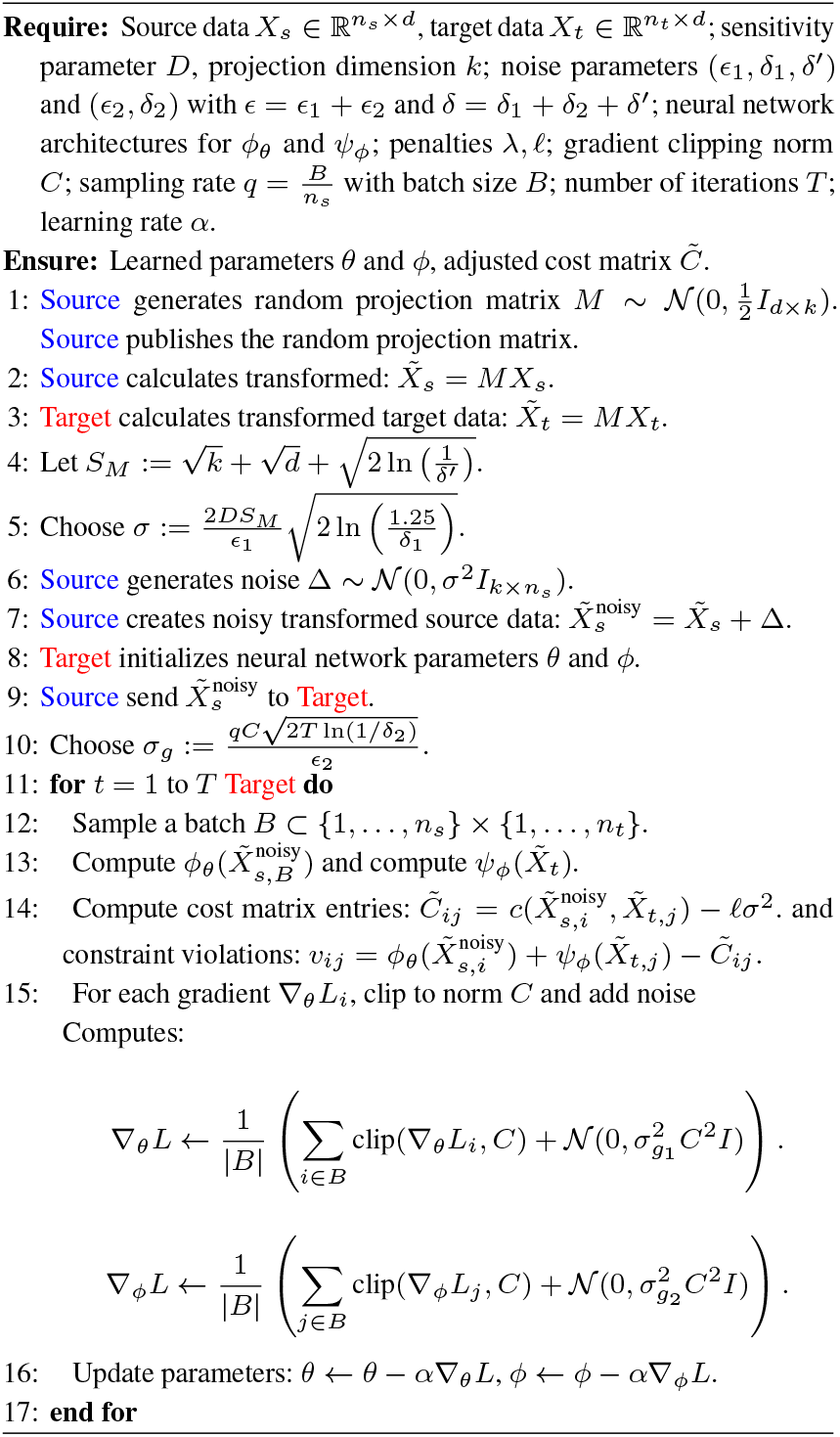

### 4.2 Implementation Considerations

In practice, the choice of the neural network architectures for *ϕ*_*θ*_ and *ψ*_*ϕ*_ plays a crucial role in the success of the method. Typically, fully connected networks with non-linear activations such as ReLU are used, with sufficient depth and width to capture the complexity of the data. The penalty parameter *λ* must be chosen carefully; a large value enforces the dual constraints more strictly but may slow down convergence, whereas a small value can lead to constraint violations. Additionally, batch processing can be employed to handle large datasets efficiently, and advanced optimization algorithms such as Adam may be used to improve convergence.

## 5 Theoretical Results

### 5.1 Privacy Guarantees for NoisyFlow

#### Theorem 5.1

(Differential Privacy w.r.t. the **Source** Dataset). *Let* 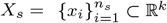 *be the source dataset, where each x*_*i*_ *satisfies* ||*x*_*i*_||_2_ ≤ *D for some D >* 0. *Let M* ∈ ℝ^*k×t*^ *be a random matrix with entries drawn independently from N* (0, 1), *and let* 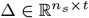 *be a noise matrix with entries drawn independently from* 𝒩 (0, *σ*^2^). *Define the perturbed dataset* 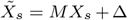. *Assume that the algorithm trains a model using differentially private stochastic gradient descent (DP-SGD) with noise parameter σ*_*g*1_ *and gradient clipping norm C. Then, for any ϵ*, *δ >* 0, *the optimal transport plan is* (*ϵ*, *δ*)*-differentially private with respect to X*_*s*_, *provided that:*

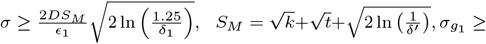

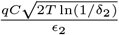, *where* 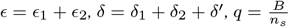 *is the sampling rate with batch size B, T is the number of iterations, and δ*^*l*^ *is a small failure probability in bounding the spectral norm of M*.

#### Theorem 5.2

(Differential Privacy w.r.t. the **Target** Dataset). *Let* 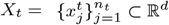 *be the target dataset with* 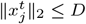.

*Then running NoisyFlow on* 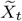 *with noise parameter σ* _*g*2_ *and gradient clipping C, the optimal transport plan are* (*ε*, *δ*)*-DP with respect to the* target *dataset, if* 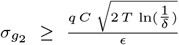 *with* 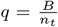 *the mini-batch sampling rate, T the total number of training iterations, and B the batch size*.

As established in Theorem 5.1 (and its counterpart for target data), NoisyFlow achieves (*ε*, δ)-differential privacy by combining noise injection with a differentially private training procedure. Specifically, two layers of noise control are employed: First, *Pre-Projection Noise* on the source data, scaled by *σ* to mask each individual point after Johnson-Lindenstrauss projection. Second, *DP-SGD Noise* in gradient computation, ensuring that model updates do not leak private information about any single participant. These steps jointly bound the privacy loss *ϵ* within the specified budget, even when an adversary has auxiliary knowledge. Consequently, membership inference attacks or sensitive attribute reconstructions are thwarted with high probability.

### 5.2 Utility Guarantees for NoisyFlow

Here, we presents a complete analysis of the approximation error incurred by the proposed differentially private optimal transport method.

#### Theorem 5.3.

*Let X*_*s*_, *X*_*t*_ ⊂ ℝ ^*d*^ *and define* 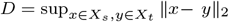. *Consider the Wasserstein distance W* (*X*_*s*_, *X*_*t*_) *with cost* 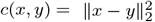. *Let M be a suitable Gaussian JL projection with parameter η, and let* 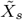, *MX*_*t*_ *be its embeddings. Introduce Gaussian noise* Δ *to form* 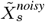, *and define an unbiased adjusted cost by subtracting kσ*^2^. *Approximate the optimal dual potentials by neural networks ϕ*_*θ*_, *ψ*_*ϕ*_ *with uniform approximation errors ϵ*_*ϕ*_, *ϵ*_*ψ*_.

*There exist constants C*_1_, *C*_2_ *>* 0 *such that with high probability*,

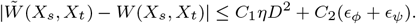

Theorem 5.3 suggests that the overall approximation error in the Wasserstein distance—after applying projection, noise addition, and neural-network approximation—remains bounded by terms that scale with the dimensionality-reduction distortion (*ηD*^2^) and the approximation error of the dual potentials (*ϵ*_*ϕ*_ + *ϵ*_*ψ*_). In practice, this indicates that NoisyFlow can maintain high fidelity in learned transport plans, thereby preserving essential structure in high-dimensional biomedical data. As a result, downstream tasks (e.g., disease classification or biomarker discovery) can maintain accuracy comparable to a non-private baseline, reflecting an advantageous trade-off between data utility and privacy protection.

Next, we incorporate a practical application scenario to illustrate the framework’s theoretical guarantee in a real-world biomedical data-sharing context. Consider a data-sharing setup involving two hospitals: Hospital A and Hospital B. Hospital A serves as the source institution, holding a dataset 𝒟_*s*_ of *n*_*s*_ patient records, each containing sensitive biomedical features (e.g., single-cell RNA-seq profiles) and possibly clinical labels (e.g., disease status). Hospital B acts as the target institution, holding a separate dataset 𝒟_*t*_ of *n*_*t*_ patient records, which may be unlabeled or sparsely labeled. The goal is to align the distributions of 𝒟_*s*_ and 𝒟_*t*_ using NoisyFlow to enable knowledge transfer without compromising patient privacy.

To formalize the privacy guarantees, let Patient *P* denote a specific individual in Hospital A’s dataset 𝒟_*s*_ whose privacy an adversary aims to compromise. The NoisyFlow framework incorporates differential privacy mechanisms to protect against membership inference and attribute inference.

#### Membership Privacy

Membership privacy guarantees that whether a particular patient *P* is included in the source dataset 𝒟_*s*_ remains indistinguishable to an adversary, even if they have access to the final model parameters or the aligned data embeddings. Specifically, consider two nearly identical datasets: 𝒟_*s*_ and 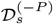, which differ only by the inclusion or exclusion of patient *P*. In the NoisyFlow framework, differential privacy (DP) ensures that an adversary at Hospital B or from a third-party cannot reliably determine whether *P*’s data is part of 𝒟_*s*_. This protection is achieved through the application of DP mechanisms that obscure the presence of any single individual’s data, thereby safeguarding membership privacy. Notice that NoisyFlow provides the same protection for Hospital A.

#### Attribute Privacy

Attribute privacy ensures that sensitive information about a patient *P* (such as disease status or specific genetic markers) cannot be inferred from the outputs generated by NoisyFlow, even if an adversary knows that *P* is included in the source dataset 𝒟_*s*_. This is particularly crucial in biomedical contexts where attributes like disease labels or rare genetic variants pose significant risks if misused. In NoisyFlow, attribute privacy is maintained by introducing noise at both the data preprocessing and model training stages. Specifically, Gaussian noise is added to the source data 𝒟_*s*_, disrupting direct associations between sensitive attributes and the resulting noisy embeddings 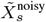. Additionally, differentially private stochastic gradient descent (DP-SGD) introduces noise during each gradient update and limits the influence of any single data point through gradient clipping. These combined measures ensure that sensitive attributes of patient *P* are not encoded in the final transport map or the aligned distributions, thereby keeping the adversary’s ability to infer *P*’s attributes highly uncertain.

## 6 Empirical results

### 6.1 Application in Functional Genomics

#### Datasets Description

The dataset used is sourced from Emani *et al*. (2024) brainSCOPE (brain Single-Cell Omics for PsychENCODE). (Emani *et al*., 2024) dataset, which compiles multi-omic single-cell data from 388 human prefrontal cortex (PFC) samples. The cohort includes individuals who are diverse in terms of biological sex, ancestry, and age, as well as healthy controls and individuals with neurological disorders. Specifically, it includes 182 controls, and individuals diagnosed with schizophrenia (*n* = 77), bipolar disorder (*n* = 34), autism spectrum disorder (*n* = 52), Alzheimer’s disease (*n* = 33), and a few with MDD/PTSD (*n* = 10). The dataset is composed of more than 2.8 million nuclei obtained from snRNA-seq, snATAC-seq, and snMultiome technologies, after stringent quality control filtering from an initial pool of nearly 4 million nuclei. Data for the cohort was harmonized from multiple external and PsychENCODE studies to provide a unified resource. This included generating uniform genotypes from whole-genome sequencing, SNP arrays, and snRNA-seq data, enabling cross-study analyses and integration.

Given the sensitive nature of health data, especially with psychiatric and neurological disorders like schizophrenia and Alzheimer’s disease, ensuring patient confidentiality while allowing meaningful data analysis is critical. By applying differential privacy, we aim to protect individual-level information in the dataset while still enabling the identification of significant cell-type-specific patterns and gene expression changes that could drive future research in neuroscience.

#### Dataset Preprocessing

To prepare the single-cell dataset for the classification task, we implemented a robust preprocessing pipeline designed to aggregate, filter, and reduce the data while preserving biologically meaningful variation. This pipeline begins with pseudobulking, which single-cell data is aggregated into cell-type-specific profiles to mitigate noise inherent in single-cell measurements. Specifically, cells were grouped by their annotated cell types, including L23IT, Oligo, Astro, L4.IT, OPC and more, which are prominent in the prefrontal cortex. For each cell type within a given patient, gene expression counts were summed to create a pseudobulk profile. To reduce dimensionality and focus the analysis on biologically relevant signals, we identified the top 2,000 highly variable genes (HVGs) using a variance-based method. HVGs were computed within each cell type across all samples to capture genes exhibiting the greatest variability, as they are often associated with key biological processes and differences. By selecting only these most informative genes, the dataset was filtered to retain features likely to contribute meaningfully to the classification task while minimizing irrelevant noise. Following pseudobulking and HVG selection, the dataset underwent feature extraction using principal component analysis (PCA). PCA was applied to the log-normalized pseudobulk profiles, reducing the dimensionality of the data while retaining the most significant axes of variation. This step compressed the data into the top principal components, which together captured the majority of the biologically relevant variance in the gene expression profiles. These principal components were then used as input features for the classification model. The final input matrix for classification was constructed by concatenating features derived from all considered cell types. The goal of the classification task was to predict whether a patient had a neurodegenerative disorder, such as Alzheimer’s disease or schizophrenia.

#### Baselines and metrics

To evaluate the efficacy of our differentially private optimal transport algorithms, we designed a study inspired by the framework outlined in the Figure 1(**b**). The dataset was split into two parts: patients belonging to the CMC cohort, representing the source institution (*n* = 70), and patients from other cohorts aside from CMC, representing the target institution (*n* = 270). The target dataset requires predictions on whether patients have a neurodegenerative disorder, and the challenge lies in aligning its distribution with that of the source dataset without violating privacy constraints. As direct data sharing between the source and target institutions is disallowed.

To assess the quality of this alignment, we measure the Wasserstein distance, which quantifies the degree of distribution alignment between the source and target datasets. Additionally, we evaluate the remapped source data using the target institution’s pre-trained XGBoost model and report two key performance metrics: the F1-score and classification accuracy, which reflects overall predictive performance.

Figure 2 illustrates key trends in the performance of our approach across various privacy-preserving and adaptation scenarios. The baseline configuration, which applies no transport to the target data, shows poor performance and a high initial Wasserstein distance, indicating a pronounced distribution mismatch between the source and target datasets.

**Fig. 2.**
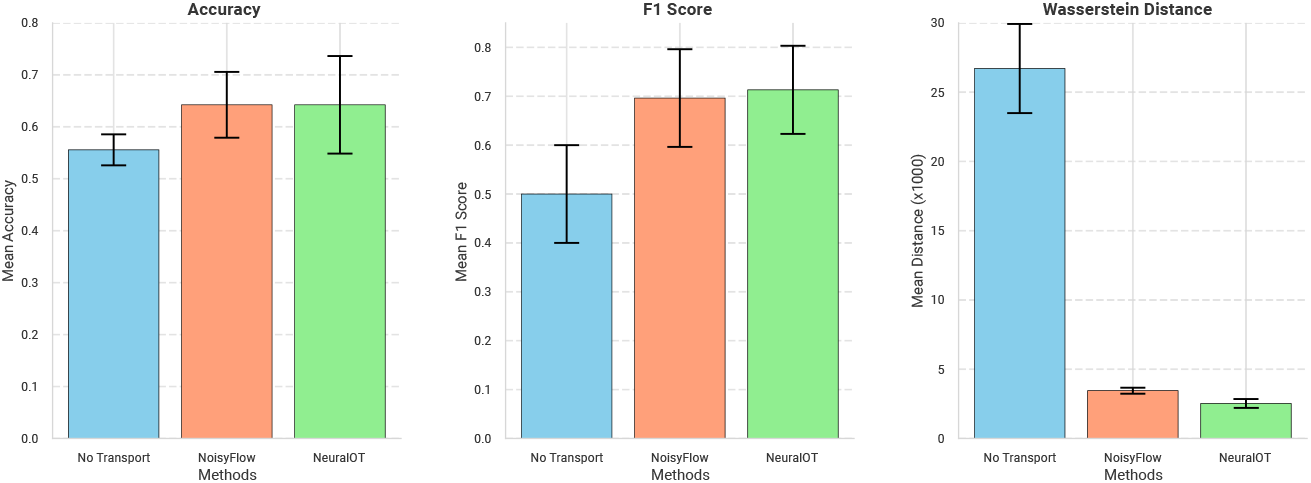
Results on the BrainSCOPE dataset are shown. Blue indicates no transport is applied, resulting in poor performance, which highlights a distribution gap between the source and target datasets. Red represents the results for NoisyFlow. Notably, NoisyFlow performs comparably to its non-DP counterparts (Green) while providing significant privacy protection for both the source and target datasets.

In contrast, NoisyFlow (red) not only demonstrates performance on par with non-differentially private counterparts (green) but also achieves a significantly lower Wasserstein distance compared to the no-transformation baseline. Furthermore, the Wasserstein distance of NoisyFlow approaches that of its non-private counterparts, highlighting its high utlity. These results underscore NoisyFlow’s potential as a viable alternative to traditional optimal transport methods, achieving effective distribution alignment while providing robust privacy guarantees.

### 6.2 Application in Biomedical Imaging

#### Dataset Description

The CAMELYON17 (Koh *et al*., 2021) dataset comprises an extensive collection of whole-slide histopathology images (WSIs) from sentinel lymph nodes of breast cancer patients. Developed for the CAMELYON17 challenge, it aims to advance automated methods for detecting and classifying metastases within lymph node tissues. Unlike its predecessor, CAMELYON16, CAMELYON17 includes slides from multiple medical centers, encompassing a variety of staining protocols and scanner settings.

Each WSI is stained with Hematoxylin and Eosin (H&E) and annotated by experts to delineate metastatic regions, ranging from small micrometastases to large macrometastases. This combination of diverse sources and precise annotations supports the development of robust and generalizable computer-aided diagnostic tools.

#### Experimental Setups

To assess NoisyFlow, we utilized source datasets comprising images from two hospitals and a target dataset containing images from three hospitals. The objective was to apply the target dataset’s model to make predictions on the source data without compromising patient privacy. To achieve this, we employed NoisyFlow to transfer data from the source to the target dataset, ensuring privacy preservation while aligning distributions to facilitate effective application of the target model. We evaluated three methods: (1) using the source dataset directly without transport, (2) applying neural optimal transport followed by re-training with the target institution’s model, and (3) utilizing NoisyFlow to securely transfer data before re-training with the target institution’s model.

**Table 1.** Performance Comparison of Data Transfer Approaches on CAMELYON17.

| Approach | Accuracy (%) | F1 (%) | Privacy |
| --- | --- | --- | --- |
| Direct Use without Transport | 75.76% | 76.23% | No |
| Neural Optimal Transport + Re-training | 86.60% | 83.70% | No |
| **NoisyFlow + Re-training** | **84.70%** | **82.90%** | **Yes** |

In our evaluation on the CAMELYON17 biomedical imaging dataset, NoisyFlow demonstrated robust performance by achieving an accuracy of 84.7% and an F1 score of 82.9%, which are comparable to the neural optimal transport approach (86.6% accuracy and 83.7% F1) while providing differential privacy guarantees. Unlike the direct use of source data without transport, which yielded significantly lower metrics (75.76% accuracy and 76.2% F1), NoisyFlow effectively aligned the distributions across multiple hospitals, addressing the distribution mismatch inherent in diverse medical datasets. This balance between maintaining high diagnostic performance and ensuring patient privacy underscores NoisyFlow’s viability as a privacy-preserving alternative to traditional optimal transport methods, making it a promising solution for deploying machine learning models in sensitive biomedical contexts.

### 6.3 Limitations and Future Work

Despite its promising capabilities, NoisyFlow presents several limitations. The empirical evaluations, although indicative of the framework’s effectiveness, were conducted on relatively small and specific datasets. Specifically, the BrainSCOPE and CAMELYON17 datasets, while representative, may not capture the full diversity and complexity of real-world biomedical data. Additionally, the scalability of NoisyFlow to larger datasets and higher-dimensional feature spaces remains an open question. The computational overhead introduced by neural network training and differential privacy mechanisms, such as DP-SGD, could pose challenges in resource-constrained settings. Furthermore, the current framework assumes that source and target domains can be effectively aligned using optimal transport, which may not hold in scenarios with extreme distribution discrepancies or non-overlapping feature spaces.

Building upon NoisyFlow, future research can explore several avenues. First, expanding empirical evaluations to include a broader range of biomedical datasets—spanning different modalities such as proteomics, metabolomics, and medical imaging—will help validate the generalizability of the framework. Second, investigating advanced neural network architectures, such as transformer-based models or state-space-models (SSM), could enhance the flexibility and expressiveness of the dual potentials, potentially improving both alignment accuracy and computational efficiency. Third, integrating adaptive differential privacy techniques that dynamically adjust privacy parameters based on data characteristics may offer better utility-privacy trade-offs. Lastly, exploring federated learning paradigms in conjunction with NoisyFlow could further decentralize the data alignment process, reducing the need for centralized data handling and enhancing privacy.

## 7 Conclusion

In summary, NoisyFlow introduces an integration of differential privacy and optimal transport, specifically designed to address the complex requirements of biomedical data sharing. By protecting patient privacy while enabling effective distribution alignment, it promises to enhance collaborative and inclusive biomedical research initiatives. Continued refinement and extensive validation will solidify its role as a foundational tool for the secure sharing and analysis of sensitive biomedical data.

## Supporting information

Proof of the theorems

## References

Alhamoud, K. et al. (2024). Fedmedicl: Towards holistic evaluation of distribution shifts in federated medical imaging. In International Conference on Medical Image Computing and Computer-Assisted Intervention, pages 383–393. Springer.

Alrefaei, A. F. et al. (2022). Genetic data sharing and artificial intelligence in the era of personalized medicine based on a cross-sectional analysis of the saudi human genome program. Scientific Reports, 12(1).

Amorim, M. et al. (2022). Benefits and risks of sharing genomic data for research: Comparing the views of rare disease patients, informal carers and healthcare professionals. International Journal of Environmental Research and Public Health, 19(14), 8788.

Bonomi, L. et al. (2020). Privacy challenges and research opportunities for genomic data sharing. Nature Genetics, 52(7), 646–654.

Bunne, C. et al. (2023). Learning single-cell perturbation responses using neural optimal transport. Nature Methods, 20(11), 1759–1768.

Cao, T. et al. (2021). Differentially private generative models through optimal transport.

Chen, J. et al. (2020). Differential privacy protection against membership inference attack on machine learning for genomic data. In Biocomputing 2021. WORLD SCIENTIFIC.

Clayton, E. W. et al. (2019). The law of genetic privacy: applications, implications, and limitations. Journal of Law and the Biosciences, 6(1), 1–36.

Emani, P. S. et al. (2024). Single-cell genomics and regulatory networks for 388 human brains. bioRxiv. Preprint.

Gentry, C. (2009). Fully homomorphic encryption using ideal lattices. In Proceedings of the 41st Annual ACM Symposium on Theory of Computing (STOC ‘09). ACM Press.

Gürsoy, G. et al. (2022). Functional genomics data: privacy risk assessment and technological mitigation. Nature Reviews Genetics, 23(4), 245–258.

Gürsoy, G. et al. (2020). Data sanitization to reduce private information leakage from functional genomics. Cell, 183(4), 905–917.e16.

Harmanci, A. O. and Gerstein, M. (2016). Quantification of private information leakage from phenotype-genotype data: linking attacks. Nature Methods, 13, 251–256.

He, Z. et al. (2018). Achieving differential privacy of genomic data releasing via belief propagation. Tsinghua Science and Technology, 23, 389–395.

Joodaki, M. et al. (2023). Detection of patient-level distances from single cell genomics and pathomics data with optimal transport (pilot). Molecular Systems Biology, 20(2), 57–74.

Jwa, A. S. and Poldrack, R. A. (2022). Addressing privacy risk in neuroscience data: from data protection to harm prevention. Journal of Law and the Biosciences, 9(2), sac025.

Koh, P. W. et al. (2021). Wilds: A benchmark of in-the-wild distribution shifts. In International conference on machine learning, pages 5637–5664. PMLR.

LeTien, N. et al. (2019). Differentially private optimal transport: Application to domain adaptation. In Proceedings of the Twenty-Eighth International Joint Conference on Artificial Intelligence, IJCAI-19, pages 2852–2858. International Joint Conferences on Artificial Intelligence Organization.

Litjens, G. et al. (2018). 1399 h&e-stained sentinel lymph node sections of breast cancer patients: the camelyon dataset. GigaScience, 7(6).

M., V. M. (2019). Melanoma skin cancer detection using image processing and machine learning. International Journal of Trend in Scientific Research and Development (IJTSRD), 3(4), 780–784.

Packhäuser, K. et al. (2022). Deep learning-based patient re-identification is able to exploit the biometric nature of medical chest x-ray data. Scientific Reports, 12, 14851.

Pajuste, F. D. et al. (2017). Fastgt: an alignment-free method for calling common snvs directly from raw sequencing reads. Scientific Reports, 7, 2537.

Panzade, P. et al. (2024). Medblindtuner: Towards privacy-preserving fine-tuning on biomedical images with transformers and fully homomorphic encryption. arXiv preprint 2401.09604.

Roberts, J. S. et al. (2020). Genetic testing for neurodegenerative diseases: Ethical and health communication challenges. Neurobiology of Disease, 141, 104871.

Schadt, E. et al. (2012). Bayesian method to predict individual snp genotypes from gene expression data. Nature Genetics, 44(5), 603–608.

Sha, Y. et al. (2023). Reconstructing growth and dynamic trajectories from single-cell transcriptomics data. Nature Machine Intelligence, 6(1), 25–39.

Song, J. et al. (2024). Privacy-preserving identification of cancer subtype-specific driver genes based on multigenomics data with privatedriver. Journal of Computational Biology, 31(2), 99–116. PMID: 38271572.

Wan, Z. et al. (2022). Sociotechnical safeguards for genomic data privacy. Nature Reviews Genetics, 23(7), 429–445.

Wang, M. et al. (2023). Identifying personal physiological data risks to the internet of everything: the case of facial data breach risks. Humanities and Social Sciences Communications, 10(1).

Warnat-Herresthal, S. et al. (2021). Swarm learning for decentralized and confidential clinical machine learning. Nature, 594, 265–270.

Yilmaz, E. et al. (2021). Genomic data sharing under dependent local differential privacy.

Zhang, J. et al. (2021). A review on modern computational optimal transport methods with applications in biomedical research. Modern Statistical Methods for Health Research, pages 279–300.

Zou, J. and Schiebinger, L. (2021). Ensuring that biomedical AI benefits diverse populations. EBioMedicine, 67(103358), 103358.

