## Supplementary material for "NoisyFlow: Differentially Private Optimal Transport Using Neural Networks for Secure Biomedical Data Sharing": Proof of the theorems

January 31, 2025

### 1 Proof of Main Theorems

#### 1.1 Proof of Main Theorem 5.1

*Proof.* Let us establish the $(\epsilon, \delta)$-differential privacy guarantees through a rigorous analysis of the algorithmic mechanisms. We proceed by analyzing each component separately and then employ the composition theorem for differential privacy.

**Definition 1.1 (Preprocessing Mechanism)** *Define $\mathcal{M}_1 : \mathcal{X}^n \to \mathbb{R}^{n\times k}$ as the mechanism that maps the source dataset $X_s$ to its perturbed representation:*

$$\mathcal{M}_1(X_s) = MX_s + \Delta, \quad \text{where } M \in \mathbb{R}^{t\times k}, \Delta \in \mathbb{R}^{n\times k}$$

First, we establish a key result regarding the sensitivity of our preprocessing mechanism.

**Lemma 1.1 (Sensitivity Bound)** *For any pair of adjacent datasets $X_s, X_s' \in \mathcal{X}^n$ differing in exactly one element, with probability at least $1-\delta'$ over the random draw of $M$, the $\ell_2$-sensitivity of $\mathcal{M}_1$ satisfies:*

$$\Delta_{\mathcal{M}_1} = \sup_{X_s \sim X_s'} \|\mathcal{M}_1(X_s) - \mathcal{M}_1(X_s')\|_F \le 2D\left(\sqrt{k} + \sqrt{d} + \sqrt{2\ln(1/\delta')}\right)$$

*where $D$ bounds the $\ell_2$-norm of input vectors.*

*Proof of Lemma.* Let $X_s$ and $X_s'$ be adjacent datasets differing at index $i$. Then:

$$\begin{aligned}
\|\mathcal{M}_1(X_s) - \mathcal{M}_1(X_s')\|_F &= \|(X_s - X_s')M\|_F \\
&= \|(x_i - x_i')M\|_2 && \text{(by adjacency)} \\
&\le \|x_i - x_i'\|_2 \|M\|_2 && \text{(by submultiplicativity)} \\
&\le 2D\|M\|_2 && \text{(by bounded norm assumption)}
\end{aligned}$$

For the random Gaussian matrix $M$ with i.i.d. entries from $\mathcal{N}(0,1)$, we invoke the following concentration result:

**Theorem 1.2 (Random Matrix Concentration)** *For $M \in \mathbb{R}^{t\times k}$ with i.i.d. entries from $\mathcal{N}(0,1)$, for any $u > 0$:*

$$\mathbb{P}\left(\|M\|_2 \ge \sqrt{k} + \sqrt{d} + u\right) \le e^{-u^2/2}$$

*Proof.* Let $\sigma_1(M)$ denote the largest singular value of $M$. We proceed in steps:

First, observe that $\|M\|_2 = \sigma_1(M)$. For a Gaussian matrix $M$, its singular values follow a specific distribution. By the concentration of Lipschitz functions of Gaussian variables:

$$\mathbb{P}(\sigma_1(M) - \mathbb{E}[\sigma_1(M)] \ge u) \le e^{-u^2/2}$$

For the expected value, we have the tight bound:

$$\mathbb{E}[\sigma_1(M)] \le \sqrt{d} + \sqrt{k}$$

This follows from Gordon's theorem on Gaussian processes.

Combining these results:

$$\mathbb{P}(\|M\|_2 \geq \sqrt{d} + \sqrt{k} + u) \leq \mathbb{P}(\sigma_1(M) - \mathbb{E}[\sigma_1(M)] \geq u) \leq e^{-u^2/2}$$

Setting $u = \sqrt{2\ln(1/\delta')}$ yields the desired bound with probability $1 - \delta'$. □

We now establish the privacy guarantee for the preprocessing step.

**Proposition 1.3 (Privacy of Preprocessing)** *Under the conditions of Lemma 1, if the noise matrix $\Delta$ has entries drawn independently from $\mathcal{N}(0, \sigma^2)$ where:*

$$\sigma \geq \frac{2D(\sqrt{k} + \sqrt{d} + \sqrt{2\ln(1/\delta')})}{\epsilon_1}\sqrt{2\ln(1.25/\delta_1)}$$

*then $\mathcal{M}_1$ satisfies $(\epsilon_1, \delta_1 + \delta')$-differential privacy.*

*Proof of Proposition.* Given the sensitivity bound from Lemma 1, we apply the Gaussian mechanism theorem:

**Theorem 1.4 (Gaussian Mechanism)** *For a function $f$ with $\ell_2$-sensitivity $\Delta_f$, the mechanism $\mathcal{M}(x) = f(x) + \mathcal{N}(0, \sigma^2)$ satisfies $(\epsilon, \delta)$-differential privacy if:*

$$\sigma \geq \frac{\Delta_f}{\epsilon}\sqrt{2\ln(1.25/\delta)}$$

Substituting $\Delta_{\mathcal{M}_1}$ and accounting for the failure probability $\delta'$ in the sensitivity bound yields the result. □

For the second component, let $\mathcal{M}_2$ denote the DP-SGD training mechanism operating on the perturbed data $\tilde{X}_s$.

Here, we first justify that employing differentially private stochastic gradient descent (DP-SGD) for training the dual potential is necessary, even when the perturbed dataset $\tilde{X}_s$ is differentially private with respect to $X_s$. Without DP-SGD, the training process can compromise the overall differential privacy guarantee for $X_s$ due to potential overfitting and sensitivity of the learned parameters to individual data points.

**Lemma 1.5 (Sensitivity of Standard SGD)** *Let $\mathcal{M}_1$ produce two perturbed datasets $\tilde{X}_s = \mathcal{M}_1(X_s)$ and $\tilde{X}'_s = \mathcal{M}_1(X'_s)$ corresponding to neighboring datasets $X_s$ and $X'_s$. The sensitivity of the model parameters $\theta$ learned using standard SGD is unbounded:*

$$\|\theta - \theta'\| = \|SGD(\tilde{X}_s) - SGD(\tilde{X}'_s)\| \text{ may not be bounded.}$$

*Proof.* Training with standard SGD involves iterative updates of model parameters $\theta$ based on gradients computed over the dataset $\tilde{X}_s$. For two neighboring datasets $\tilde{X}_s$ and $\tilde{X}'_s$, the gradients may differ significantly due to the unbounded sensitivity of gradient computation:

$$g_i = \nabla_\theta L(\theta; x_i), \quad g'_i = \nabla_\theta L(\theta; x'_i).$$

Thus, the cumulative effect of these differences across all iterations can cause $\|\theta - \theta'\|$ to grow arbitrarily large, especially in the absence of gradient clipping or noise addition. □

Differentially private SGD ensures that the sensitivity of model parameters to changes in the input dataset is bounded. We now formalize the privacy guarantee of DP-SGD.

**Theorem 1.6 (Differential Privacy of DP-SGD)** *Let $\tilde{X}_s$ be a dataset, and let $\mathcal{M}_2$ denote the training mechanism using DP-SGD. If the per-example gradients are clipped to a norm $C$, and Gaussian noise with standard deviation $\sigma_g$ is added to the clipped gradients, $\mathcal{M}_2$ satisfies $(\epsilon_2, \delta_2)$-differential privacy with respect to $\tilde{X}_s$, provided that:*

$$\sigma_g \geq \frac{qC\sqrt{2T\ln(1/\delta_2)}}{\epsilon_2},$$

*where $q = B/n_s$ is the sampling rate, $T$ is the number of iterations, and $B$ is the batch size.*

Finally, we combine these results using the advanced composition theorem for differential privacy.

**Theorem 1.7 (Composition)** *For mechanisms $\{\mathcal{M}_i\}_{i=1}^k$ where each $\mathcal{M}_i$ is $(\epsilon_i, \delta_i)$-differentially private, their composition satisfies $(\sum_i \epsilon_i, \sum_i \delta_i)$-differential privacy.*

Applying this theorem to $\mathcal{M}_1$ and $\mathcal{M}_2$ with parameters:

$$\epsilon_1 = \epsilon_2 = \epsilon/2$$
$$\delta_1 = \delta_2 = \delta' = \delta/3$$

yields the overall $(\epsilon, \delta)$-differential privacy guarantee. □

**Remark.** By clipping per-sample gradients to bound their sensitivity, adding appropriately calibrated Gaussian noise to the aggregated gradients, and accounting for privacy loss over multiple iterations using the advanced composition theorem, the neural network-based optimal transport algorithm satisfies the $(\varepsilon_{\text{total}}, \delta_{\text{total}})$-definition of differential privacy. This ensures that the inclusion or exclusion of any single data point in the source dataset $X_s$ has a limited impact on the probability distribution of the algorithm's outputs, thereby protecting individual privacy.

□

#### 1.2 Proof of Main Theorem 5.2

This follows from the standard proof of DP-SGD and is omitted here.

#### 1.3 Proof of Theorem 5.3

We begin by stating the main theorem that bounds the difference between the true Wasserstein distance and the approximated one obtained after random projection, noise injection with unbiased correction, and neural approximation of the dual potentials. Subsequent theorems and lemmas are then used to prove this main result. Compared to previous statements, we now explicitly incorporate the maximum pairwise distance factor in the Johnson–Lindenstrauss (JL) approximation term and carefully present the duality theorem for optimal transport, as well as a detailed error analysis for the neural approximation of the dual potentials. The proof of Theorem **??** follows from three key components: (1) a JL lemma ensuring small distortion of squared distances, (2) a noise correction lemma showing unbiasedness, and (3) a neural approximation lemma bounding the error from imperfect dual potentials. Each of these results is presented and proved in the following sections.

**Theorem 1.8 (Johnson–Lindenstrauss with Diameter)** *Let* $X_s \cup X_t \subset \mathbb{R}^d$ *and define* $D = \sup_{x \in X_s, y \in X_t} \|x - y\|_2$. *For any* $\eta \in [0, 0.5]$, *let* $M \in \mathbb{R}^{d \times k}$ *have i.i.d. entries from* $\mathcal{N}(0, 1/\ell)$. *With probability at least* $1 - \delta_{JL}$,

$$(1-\eta)\|x_s - x_t\|_2^2 \le \|\tilde{x}_s - \tilde{x}_t\|_2^2 \le (1+\eta)\|x_s - x_t\|_2^2$$

*holds for all* $(x_s, x_t) \in X_s \times X_t$. *Consequently,*

$$|\|\tilde{x}_s - \tilde{x}_t\|_2^2 - \|x_s - x_t\|_2^2| \le \eta\|x_s - x_t\|_2^2 \le \eta D^2.$$

*Proof.* This follows from the Gaussian Johnson–Lindenstrauss lemma applied to the set $X_s \cup X_t$. The probability bound and distortion parameter $\eta$ are standard results. The inequality involving $D^2$ arises by noting that the maximum squared distance between points is at most $D^2$, so the maximal difference induced by the $(1 \pm \eta)$ factor is at most $\eta D^2$. □

**Lemma 1.9 (Unbiased Noise Correction)** *Let* $\tilde{X}_s = MX_s$ *and* $\Delta \sim \mathcal{N}(0, \sigma^2 I_{k \times n_s})$. *Define* $\tilde{X}_s^{noisy} = \tilde{X}_s + \Delta$ *and consider* $c(x, y) = \|x - y\|_2^2$. *Then*

$$\mathbb{E}_\Delta[\|\tilde{x}_s + \Delta - \tilde{x}_t\|_2^2] = \|\tilde{x}_s - \tilde{x}_t\|_2^2 + k\sigma^2.$$

*By defining* $\tilde{c}(\tilde{x}_s^{noisy}, \tilde{x}_t) = \|\tilde{x}_s^{noisy} - \tilde{x}_t\|_2^2 - k\sigma^2$, *we have*

$$\mathbb{E}_\Delta[\tilde{c}(\tilde{x}_s^{noisy}, \tilde{x}_t)] = \|\tilde{x}_s - \tilde{x}_t\|_2^2.$$

*Thus, the noise-adjusted cost is unbiased in expectation.*

*Proof.* Since $\Delta \sim \mathcal{N}(0, \sigma^2 I)$, we have $\mathbb{E}[\|\Delta\|_2^2] = k\sigma^2$. Expanding

$$\mathbb{E}_\Delta[\|\tilde{x}_s + \Delta - \tilde{x}_t\|_2^2] = \mathbb{E}_\Delta[\|\tilde{x}_s - \tilde{x}_t\|_2^2 + 2\langle \Delta, \tilde{x}_s - \tilde{x}_t \rangle + \|\Delta\|_2^2].$$

Since $\mathbb{E}[\Delta] = 0$ and $\Delta$ is independent of $\tilde{x}_s, \tilde{x}_t$, the cross-term vanishes. Therefore

$$\mathbb{E}_\Delta[\|\tilde{x}_s + \Delta - \tilde{x}_t\|_2^2] = \|\tilde{x}_s - \tilde{x}_t\|_2^2 + \mathbb{E}[\|\Delta\|_2^2] = \|\tilde{x}_s - \tilde{x}_t\|_2^2 + k\sigma^2.$$

Subtracting $k\sigma^2$ yields the unbiased corrected cost. □

**Lemma 1.10 (Neural Approximation of the Dual Potentials)** *Assume that* $(\phi^*, \psi^*)$ *are optimal dual potentials satisfying*

$$W_c(\mu_s, \mu_t) = \mathbb{E}_{x_s \sim \mu_s}[\phi^*(x_s)] + \mathbb{E}_{x_t \sim \mu_t}[\psi^*(x_t)]$$

*with* $\phi^*(x_s) + \psi^*(x_t) \le c(x_s, x_t)$ *for all* $(x_s, x_t)$. *Suppose* $\phi_\theta, \psi_\phi$ *satisfy* $\sup_x |\phi_\theta(x) - \phi^*(x)| \le \epsilon_\phi$ *and* $\sup_x |\psi_\phi(x) - \psi^*(x)| \le \epsilon_\psi$. *Then*

$$|\mathbb{E}_{x_s \sim \mu_s}[\phi_\theta(x_s)] + \mathbb{E}_{x_t \sim \mu_t}[\psi_\phi(x_t)] - W_c(\mu_s, \mu_t)| \le \epsilon_\phi + \epsilon_\psi.$$

*Proof.* Let $Z_{\phi,\psi} = \mathbb{E}_{x_s \sim \mu_s}[\phi(x_s)] + \mathbb{E}_{x_t \sim \mu_t}[\psi(x_t)]$. By optimality of $\phi^*, \psi^*$, $Z_{\phi^*,\psi^*} = W_c(\mu_s, \mu_t)$. Consider $Z_{\phi_\theta,\psi_\phi}$. Since $|\phi_\theta(x) - \phi^*(x)| \le \epsilon_\phi$ and $|\psi_\phi(x) - \psi^*(x)| \le \epsilon_\psi$ for all $x$, it follows that

$$Z_{\phi_\theta,\psi_\phi} = \mathbb{E}_{x_s}[\phi_\theta(x_s)] + \mathbb{E}_{x_t}[\psi_\phi(x_t)]$$

can differ from $Z_{\phi^*,\psi^*}$ by at most $\epsilon_\phi + \epsilon_\psi$, since both expectations are over finite sets with uniform weights. Thus

$$|Z_{\phi_\theta,\psi_\phi} - Z_{\phi^*,\psi^*}| \le \epsilon_\phi + \epsilon_\psi.$$

Since $Z_{\phi^*,\psi^*} = W_c(\mu_s, \mu_t)$, the claim follows. □

*Proof of Theorem 5.3.* Using Theorem 1.8, with probability at least $1 - \delta_{JL}$, all squared distances are approximated to within a factor $(1 \pm \eta)$, and therefore differ by at most $\eta D^2$. This implies that the Wasserstein distance computed using $\tilde{X}_s, \tilde{X}_t$ without noise and dual approximations differs from $W(X_s, X_t)$ by a term on the order of $\eta D^2$. Incorporating the constant $C_1$, this difference can be bounded as $C_1 \eta D^2$.

Next, by Lemma 1.9, the introduction of noise and the subtraction of $k\sigma^2$ yields an unbiased estimator of the squared distances in expectation. Thus, no systematic bias is introduced by the noise when taking expectation. Any finite-sample variance effects vanish as $n_s, n_t$ grow large or can be absorbed into the constants. Since the problem focuses on expectation and not sample variance, no additional deterministic bias term arises from the noise.

Finally, by Lemma 1.10, the neural approximation of the dual potentials introduces an additive error at most $\epsilon_\phi + \epsilon_\psi$. Incorporating this into a constant $C_2$, we have an additional $C_2(\epsilon_\phi + \epsilon_\psi)$ error in the final approximation.

Combining these results, we obtain

$$|\tilde{W}(X_s, X_t) - W(X_s, X_t)| \le C_1 \eta D^2 + C_2(\epsilon_\phi + \epsilon_\psi)$$

as required. This proves Theorem 5.3. □

#### 1.4 Runtime Complexity

The runtime complexity of the Neural Optimal Transport with Noise Cost Adjustment algorithm under Differential Privacy is partitioned into preprocessing and iterative training phases.

**Preprocessing Phase:** Generating the random projection matrix $M \in \mathbb{R}^{d \times k}$ requires $O(dk)$ time. Transforming the source and target datasets through matrix multiplication incurs a computational cost of $O\big((n_s + n_t)dk\big)$, where $n_s$ and $n_t$ denote the number of source and target samples, respectively. Adding noise to the source data introduces an additional complexity of $O(n_s k)$, while initializing the neural network parameters entails $O(p)$, where $p$ represents the total number of parameters in the networks. Consequently, the total preprocessing cost is:

$$O\left((n_s + n_t)dk + n_s k + p\right)$$

For large-scale datasets, the term $(n_s + n_t)dk$ typically dominates, allowing the preprocessing complexity to be approximated as:

$$O\left((n_s + n_t)dk\right)$$

**Iterative Training Phase:** The training loop executes for $T$ iterations with a batch size of $B$. In each iteration, sampling a batch incurs $O(B)$ time. Computing the neural network outputs involves forward passes through networks $\phi$ and $\psi$, with complexities $O(f_\phi(B,k))$ and $O(f_\psi(n_t,k))$, respectively. The computation of the cost matrix for all batch-target pairs contributes $O(Bn_tk)$, while evaluating constraint violations and the loss function adds $O(Bn_t)$. Gradient computation, clipping, and noise addition collectively require $O(f_\phi(B,k) + f_\psi(n_t,k) + p)$, and updating the network parameters incurs an additional $O(p)$. Therefore, the per-iteration complexity is:

$$O\left(B + f_\phi(B,k) + f_\psi(n_t,k) + Bn_tk + p\right)$$

Aggregating over $T$ iterations, the total training cost becomes:

$$O\left(T \cdot \left(B + f_\phi(B,k) + f_\psi(n_t,k) + Bn_tk + p\right)\right)$$

In scenarios where the complexities $f_\phi$ and $f_\psi$, as well as the parameter size $p$, are not excessively large relative to $Bn_tk$, the term $Bn_tk$ dominates. Consequently, the training complexity simplifies to:

$$O\left(T \cdot Bn_tk\right)$$

**Overall Runtime Complexity:** Combining both preprocessing and training phases, the overall runtime complexity of the algorithm is:

$$O\left((n_s + n_t)dk + T \cdot Bn_tk\right)$$
